# An evolutionary innovation is contingent on maintaining adaptive potential until competition subsides

**DOI:** 10.1101/249110

**Authors:** Dacia Leon, Simon D'Alton, Erik M. Quandt, Jeffrey E. Barrick

## Abstract

After 15 years of the Lenski experiment one of twelve *Escherichia coli* populations evolved the ability to utilize an abundant but previously untapped carbon source, citrate. Mutations responsible for the appearance of rudimentary citrate utilization (Cit^+^ phenotype) and for refining this ability have been characterized. However, the complete nature of the genetic and/or ecological events that set the stage for this key innovation remain unknown. We found that there was a slight fitness benefit for introducing an activated *citT* cassette that mimics the mutation causing Cit^+^ into the ancestor of the evolution experiment and strains isolated from the population close to when it evolved. However, there was no benefit or even a large deleterious effect in intermediate strains. We conclude that achieving Cit^+^ was contingent on both an evolutionary trajectory that maintained a potentiated genetic state and the slowing rate of adaptation in this population late in the experiment.

## Introduction

*Escherichia coli* can more effectively uptake iron when a small amount of citrate, about 10 μM, is added to chemically defined media^1^. In 1950, when DM (Davis-Mingioli) medium was initially designed, the recipe used 1700 μM citrate. DM medium was formulated to isolate auxotrophic mutants by the penicillin method. It was shown that the addition of citrate improved the lethality of penicillin, thereby reducing the number of false-positives when selecting for auxotrophs. The penicillin method was widely used, and DM was adopted as a chemically defined medium for other types of experiments as a consequence^2^. The elevated concentration of citrate was typically unnecessary in these new circumstances, but it remained unaltered as new labs and generations of scientists inherited the same DM recipe.

The Lenski long-term evolution experiment (LTEE) consists of twelve *E. coli* B populations that have been propagated daily in glucose-limited DM medium for over 60,000 generations^3,4^. A relatively low concentration of glucose (139 μM) was used in the LTEE to restrict the cell density and thereby reduce the chances that stable ecology reinforced by cross-feeding interactions would evolve, which has largely been the case except for in one population^5,6^. The low-glucose formulation of DM used in the LTEE means that the standard amount of citrate present (1700 μM) represents a substantial nutrient pool that could be exploited, but the ancestral strain of *E. coli* is unable to utilize citrate under the conditions of the LTEE. Citrate is an untapped niche.

After 31,500 generations, a mutant in one of the LTEE populations, designated Ara-3, evolved the ability to utilize citrate as a carbon source. Descendants of this mutant that were able to fully exploit this additional carbon source evolved by 33,000 generations and dominated thereafter^7^. Previous work has demonstrated that the evolution of citrate utilization in the LTEE proceeded through three stages typical of any key innovation: potentiation, actualization, and refinement^7–10^. The principal mutations involved in the latter two steps have been characterized. Actualization refers to the first manifestation of a rudimentary Cit^+^ phenotype. The actualizing mutation is a tandem duplication of the *rnk-citG* region of the *E. coli* chromosome that includes the citrate:succinate antiporter gene, *citT*. This duplication results in an arrangement in which one of the two copies of *citT* is now downstream of the aerobically-active *rnk* promoter (P_rnk_). Thus, the actualization step results in CitT production under the LTEE conditions^8^.

However, early Cit^+^ cells with the *citT* duplication are able to uptake and metabolize only a small fraction of the citrate present during one 24-hour growth cycle of the LTEE^7,11^. Subsequently, refinement mutations improved the rudimentary Cit^+^ trait in their descendants such that they became capable of utilizing all of the citrate in DM during each growth cycle (Cit^++^ phenotype). One critical mutation for refinement activated expression of *dctA*, a C_4_-dicarboxylate:H^+^ symporter gene. DctA allows active transport of C_4_-dicarboxylates, including succinate, into the cell. Because CitT is an antiporter that couples export of these compounds to citrate import, expression of DctA creates a sustainable cycle for importing citrate that is powered by the proton gradient^9^. The *dctA* mutation refines the rudimentary Cit^+^ trait into the Cit^++^ phenotype that was responsible for the population expansion observed at ~33,000 generations in the LTEE^7^.

While the actualization and refinement stages of Cit^+^ evolution in the LTEE are understood, the mechanistic basis of potentiation has remained elusive. The critical diagnostic characteristic of a potentiated strain is that is has an increased chance of giving rise to a Cit^+^ descendant after further evolution^8^. Blount *et al*. identified potentiated strains by performing ‘replay’ experiments^7^. In these experiments, pre-Cit^+^ clones isolated from the LTEE population at various ime points were tested to determine whether they were capable of evolving citrate utilization. Cit^+^ cells rarely arose in these replay experiments. When they did, the Cit^+^ trait re-evolved more often in clones selected from later time points that were closer to when the *citT* duplication first arose in the LTEE population.

The phylogenetic distribution of the LTEE strains giving rise to Cit^+^ variants in the replay experiments suggests the existence of at least two critical junctures at which the potential for evolving Cit^+^ increased^8^. By 20,000 generations, the LTEE population had diversified into three long-lived clades that co-existed at least until full citrate utilization (Cit^++^) evolved at ~33,000 generations. *E. coli* isolates from all of these groups evolved Cit^+^ in the replay experiments, whereas no strains from earlier than 20,000 generations did, suggesting that all three clades share some determinant of potentiation. A significantly higher proportion of clones from the clade that gave rise to citrate utilization in the LTEE were able to evolve Cit^+^ in the replays, suggesting that they share a second determinant for increased potentiation not present in the other clades. Due to the extreme rarity of Cit^+^ arising in these replay experiments even after months of evolution, it is not realistic to use this approach to further narrow down the genetic basis of potentiation.

In this study, we tested the viability of the critical actualizing mutation for Cit^+^ evolution in a series of pre-Cit^+^ isolates from the LTEE by measuring the effect of activating *citT* expression on competitive fitness. We found that activating *citT* expression slightly increased the fitness of the ancestral strain and some later pre-Cit^+^ clones. Unexpectedly, activating *citT* expression was highly deleterious in certain strains from intermediate time points, and we did not find any strains that benefitted significantly more from this mutation than the ancestral strain did. We conclude that potentiation for the evolution of citrate utilization in the LTEE is due to the interplay of genetic factors in specific strains and the population at large. First, adaptation had to occur via a genetic trajectory that maintained the potential for evolving Cit^+^ by a beneficial mutational step in order for the innovation to remain accessible. Second, the rate of adaptation in the overall population needed to slow to a pace at which early variants with the weakly beneficial Cit^+^ trait could avoid being driven extinct by competitors before refining mutations arose.

## Results

### Cit^+^ was only slightly beneficial when it evolved in the LTEE

By definition, the first *E. coli* cell that evolved the citT-activating mutation that was ultimately successful in the LTEE was fully potentiated when this mutation arose. The earliest Cit^+^ descendant of this cell that has been identified is strain ZDB564 from 31,500 generations. At this time Cit^+^ cells were still extremely rare in the population^10^, which means that it is likely that the suite of mutations in ZDB564 is identical to those in the first Cit^+^ cell, or nearly so. Previously, strain ZDB706, a Cit^‒^ revertant of ZDB564, was isolated by passaging ZDB564 on DM medium lacking citrate to allow for the spontaneous collapse of the *rnk-citG* duplication to the ancestral single-copy state that lacks a copy of the *rnk* promoter upstream of *citT* (**Fig. 1a**)^10^.

**Figure 1.**
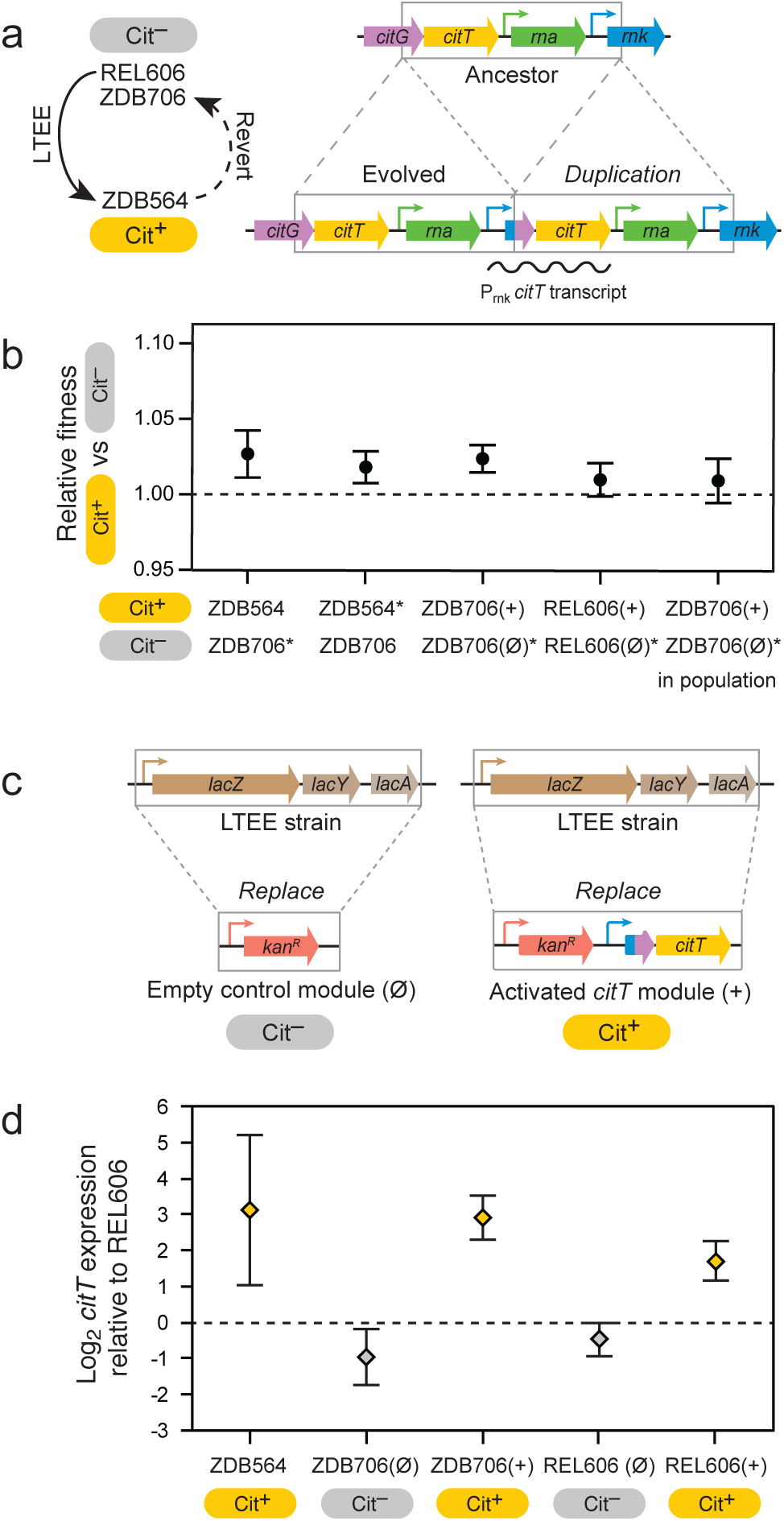
Evolution of rudimentary citrate utilization by activating *citT* expression is slightly beneficial in the genetic background in which it evolved and in the LTEE ancestor. **(a)** The *rnk-citG* duplication that evolved in the LTEE creates a genomic configuration in which a novel mRNA encoding the CitT transporter is expressed from the *rnk* promoter (P_rnk_) (right). This mutation alone is sufficient for weak citrate utilization (Cit^+^ phenotype). It is the ‘actualizing mutation’ in the evolution of this key innovation. Strain ZDB564 is the earliest Cit^+^ isolate from the LTEE. In order to measure the effect that this mutation had on competitive fitness when it evolved, a spontaneous Cit^-^ revertant of ZDB564 in which the duplication collapsed back to the ancestral state was isolated (left). **(b)** Competitive fitness of Cit^+^ versus Cit^-^ strain variants. The ZDB564 versus ZDB706 competitions measure the fitness effect of the *rnk-citG* duplication when it evolved. The ZDB706 and REL606 competitions test the effect of adding one copy of the evolved P_*rnk*_*-citT* module into a strain (^+^) versus adding an empty version of the same cassette (Ø), as pictured in **c.** An additional ZDB706 competition (in population) was conducted with the two strains together mixed at a 1:99 ratio with the evolved LTEE population from at 31,000 generations to determine if the mutation had a different effect on fitness when rare in the population. Starred strains (*) have a change to the Ara^+^ marker state to allow competition with the corresponding Ara^-^ strain as illustrated in **Figures S1** and **S2**. The marker change had no effect on competitive fitness in each case (**Fig. S3**). Error bars are 95% confidence intervals. (c) Schematic of the gene cassettes used in the P_*rnk*_*-citT* knock-in assay showing how they were integrated into the *E. coli* chromosome in a way that replaces the native *lac* locus. **(d)** *citT* mRNA expression levels measured relative to the REL606 LTEE ancestor in the evolved Cit^+^ isolate from the LTEE (ZDB564) and strains with the P_*rnk*_*-citT* and corresponding empty control cassettes integrated into their chromosomes. Error bars are 95% confidence intervals.

We co-cultured ZDB564 and ZDB706 in DM medium to estimate the effect that the *citT* duplication had on competitive fitness when it originally arose. These experiments involved reverting an arabinose-utilization allele in one of the two strains to be competed from the inactivated state present in all strains from this LTEE population (Ara^‒^) to the active state (Ara^+^) so that cells of each type can be distinguished by the colors of the colonies that they form on indicator plates (**Fig. S1** and **Methods**)^4^. These Ara^+^ strain variants were assayed to establish that the genetic marker was neutral with respect to fitness and that no secondary mutations affecting fitness had accumulated during strain construction prior to further competition experiments (**Fig. S2**). When competing ZDB564 and ZDB706, we found a slight fitness advantage of 2.2% for the presence of the citT-activating duplication in ZDB564 (**Fig. 1b**). This result that was consistent between the competitions utilizing Ara^+^ ZDB706 or Ara^+^ ZDB564 marked variants (two-tailed *t*-test, *P*=0.33, *n*=12 and 18, respectively).

### Development of P_*rnk*_*-citT* knock-in assay for potentiation

We next wanted to add the citT-activating mutation to pre-Cit^+^ strains in order to test our hypothesis that there was a transition in the lineage leading to Cit^+^ such that this mutation became more beneficial once a potentiated genetic background evolved. The effect of adding a plasmid containing the evolved P_*rnk*_*-citT* unit has been tested in previous studies,^8,9^ but this approach is problematic because these plasmids are multicopy, whereas only a single activated copy of the *citT* gene was present in the initial Cit^+^ strains. However, engineering the authentic *rnk-citG* duplication into the chromosome of a strain is difficult because this configuration is genetically unstable. It readily collapses via homologous recombination if there is not selection to maintain citrate utilization, as was utilized in reverting ZDB564 to the Cit^‒^ variant ZDB706.

To address these shortcomings, we developed a P_*rnk*_*-citT* knock-in assay, in which a mimic of the evolved configuration is integrated into the chromosome of a pre-Cit^+^ LTEE clone (Fig. 1c). Briefly, we created an activated *citT* module linked to an antibiotic selection marker in which the *rnk* promoter is upstream of the truncated *rnk-citG* fusion ORF formed by the duplication followed by the complete *citT* reading frame. To control for any fitness cost imposed by the selection marker, we also made a null module containing only the antibiotic resistance gene.

Both of these cassettes are targeted to integrate into the *E. coli* chromosome such that they replace the *lac* operon, which is unrelated to citrate or glucose metabolism.

We validated this approach by adding the P_*rnk*_*-citT* module to the fully potentiated Cit^‒^ revertant, ZDB706, and adding the null module to its neutral Ara^+^ variant. Addition of the P_rnk_-*citT* cassette to the fully potentiated Cit^‒^ strain ZDB706 resulted in increased *citT* mRNA levels equivalent to those seen in ZDB564, the original Cit^+^ isolate with the actual *rnk-citG* duplication that evolved in the LTEE (two-tailed *t*-test, *P* = 0.65, *n* = 3) (**Fig. 1d**). The resulting Cit^+^ variant of ZDB706 had a fitness advantage of 2.4% over the corresponding Cit^‒^ variant with the null knock-in cassette (**Fig. 1b**), which was not statistically different from the fitness advantage found for the authentic citT-activating mutation in the pooled ZDB564 versus ZDB706 competitions (two-tailed *t*-test, *P* = 0.71, *n* = 6 and 30, respectively). Therefore, applying the P_*rnk*_*-citT* knock-in assay to additional strains allows us to ask: if the citT-activating mutation had evolved in a genetic background that existed earlier in the LTEE, would it have been as beneficial?

### Cit^+^ would have been modestly beneficial if it evolved in the LTEE ancestor

As a first step in further elucidating the fitness consequences of evolving rudimentary Cit^+^ on other strains from the LTEE, we performed the P_*rnk*_*-citT* knock-in assay on the ancestral LTEE strain, REL606. We found a slight fitness benefit of 1.0% for the Cit^+^ mutation (**Fig. 1b**). This effect size is near the limit for the smallest differences that can be distinguished in these types of competitive fitness assays, resulting in relatively weak support for the hypothesis that there was any fitness advantage at all for the REL606 variant with the P_*rnk*_*-citT* module relative to the one with the null module (one-tailed *t*-test, *P* = 0.033, *n* = 12). There was evidence, though also not very strong, that the benefit of the P_*rnk*_*-citT* module in the fully potentiated strain ZDB706 was greater than it was in REL606 (one-tailed *t*-test, *P* = 0.018, *n* = 12 and 6, respectively). Expression of *citT* was not quite as high in the REL606 strain with the P_*rnk*_*-citT* module as it was in ZDB706 with the same module (two-tailed *t*-test, *P* = 0.00016, *n* = 3) (**Fig. 1d**), suggesting that mutations during the LTEE on the lineage leading to Cit^+^ may have altered the strength of the *rnk* promoter. Overall, the REL606 measurements indicated, surprisingly, that there was likely a modest benefit for a mutation activating expression of *citT* at the very beginning of the LTEE, and that this benefit may have only slightly improved after further mutations that occurred during the potentiation stage in the evolution of this metabolic innovation.

### No evidence for ecological potentiation

Why did the appearance of citrate utilization take so long and why has it not evolved in other LTEE populations? One hypothesis for its rarity is that the evolution of a particular ecology in the population was important for enabling the evolution of Cit^+^. This type of situation is known to occur, for example, when nutrient cross-feeding between genetically diverged subpopulations yields negative frequency dependence, such that the competitive advantage for a newly evolved strain or a certain subpopulation is greater when it is rare within the population than when it is common^12^. The pre-Cit^+^ clade was rare during the time period when the *rnk-citG* duplication evolved. It constituted < 1-5% of the population from 30,000 to 32,500 generations^10^.

To test whether this kind of ‘ecological potentiation’ was important for the evolution of Cit^+^ in the LTEE, we repeated the P_*rnk*_*-citT* knock-in assay competition for strain ZDB706 in the context of the full diversity that existed in the population at 31,000 generations (**Fig. 1b**). The Cit^+^ and Cit^‒^ variants were mixed together equally and added such that they comprised ~1% of the cells in a mixture with the evolved population sample. In this context, the Cit^+^ strain had a 0.9% fitness advantage over the Cit^‒^ strain, which was less than and only marginally different from the result when the two strains were competed versus one another normally (two-tailed *t*-test, *P* = 0.053). Thus, we find no support for the ecological potentiation hypothesis. If anything, the more diverse mixed population context may slightly reduce the benefit of Cit^+^ evolution.

### Anti-potentiated strains evolved at intermediate time points

We next performed the P_*rnk*_*-citT* knock-in assay on 23 additional clones isolated from the LTEE population (**Fig. 2a**). Our goal was to determine whether activating *citT* expression was similarly beneficial in other evolved genetic backgrounds. We measured *citT* mRNA levels in five of the constructed strains with the P_*rnk*_*-citT* cassette and found them to be similar in all of these strains (**Fig. S5**), indicating that the strength of the *rnk* promoter was largely unchanged by the specific suites of evolved mutations present in each of these strains. For four of the evolved strains we found strong evidence that the *citT* cassette significantly increased fitness versus the control with the null cassette, as it had in the fully potentiated strain ZDB706 (one-tailed bootstrap test incorporating Ara^+^/Ara^‒^ marker and Cit^+^/Cit^‒^ competitions described in **Methods**, *P* < 0.05). In nine strains, *citT* activation had no significant effect on fitness (two-tailed bootstrap test, *P* < 0.05), though our measurements did not achieve sufficient precision to rule out that there was a fitness benefit of 1% or greater in seven of these cases (one-tailed bootstrap test, *P* < 0.05).

**Figure 2.**
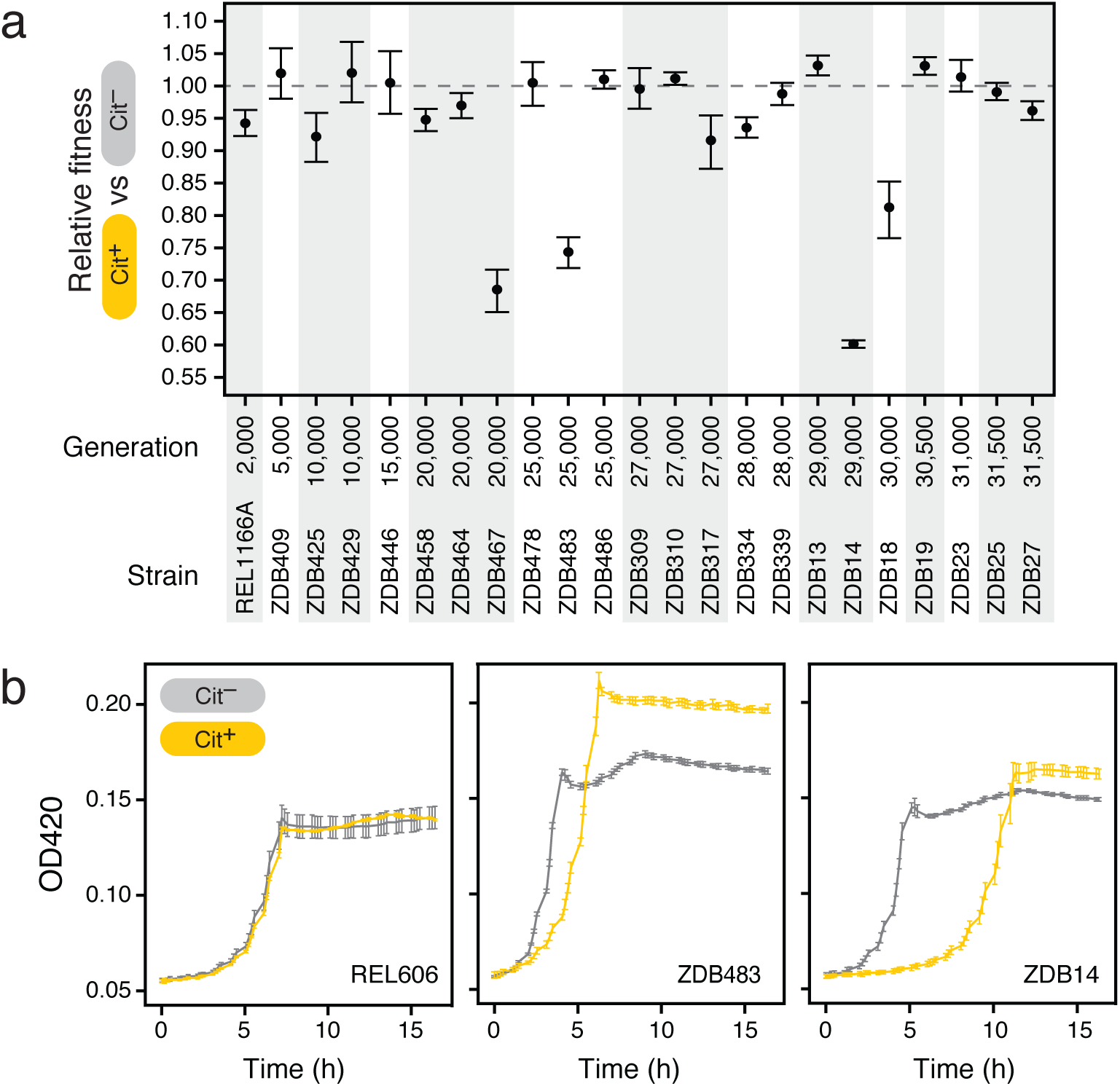
Fitness consequences of evolving Cit^+^ in different evolved genetic backgrounds. **(a)** Results of the *P*_*rn*_*k-citT* knock-in assay on 23 pre-Cit^+^ evolved strains. The clones are ordered by the generation from which they were isolated. Error bars are 95% confidence intervals. Strain construction details and how the results of competition assays were combined into these fitness estimates are described in the **Methods** and **Figures S1-S4. (b)** Increased lag phase upon addition of the P_*rnk*_*-citT* module in anti-potentiated strains. Growth curves for the ancestor, REL606, and two anti-potentiated strains, ZDB483 and ZDB14, are shown. Error bars are standard deviations of four replicate cultures.

Unexpectedly, the Cit^‒^ variant outcompeted the Cit^+^ variant for the 11 remaining strains of the 23 we tested (one-tailed bootstrap test, *P* < 0.05). The actualizing step needed for subsequently evolving full citrate utilization (Cit^++^) would have been effectively blocked if it occurred in these strain backgrounds; they are ‘anti-potentiated’. For five of these strains, activating *citT* expression was extremely detrimental, decreasing competitive fitness by >20% (one-tailed bootstrap test, *P* < 0.05). For ZDB483 and ZDB14, two of the severely anti-potentiated strains, we investigated the nature of this defect by comparing growth curves of the Cit^+^ and Cit^‒^ variants. There was very little difference in the growth curve for the LTEE ancestor REL606 whether the activated *citT* cassette or null control cassette was added to its genome, which is in keeping with its almost imperceptible effect on the competitive fitness of this strain. In contrast, we found that activating *citT* expression drastically increased the lag phase of growth in the severely anti-potentiated strains (**Fig. 2b**). This additional lag time can explain the sizable competitive disadvantage versus the Cit^‒^ strain, even though the Cit^+^ variants are able to reach a higher final cell density if cultured alone.

### Mapping potentiation onto phylogeny

Identifying specific mutations that contributed to potentiation and anti-potentiation requires interpreting the fitness data from the P_*rnk*_*-citT* knock-in assays in a phylogenetic context. To improve the resolution of a previously published whole-genome phylogenetic tree of 29 clonal isolates from this LTEE population^8^, we sequenced the genomes of 20 new clones (**Table S1**) and also incorporated 12 other clones sequenced in another recent study of the rate of genome evolution through 50,000 generations in all LTEE populations^13^. The 20 newly sequenced isolates were selected to improve our ability to temporally order mutations that occurred near when citrate utilization evolved: they were minimally diverged from the line of descent to the Cit^+^ progenitor and were mostly sampled at later time points.

The updated phylogenetic tree (**Fig. 3**) includes all 25 clones we tested with the P_*rnk*_*-citT* knock-in assay. We used these strains to identify branches in the tree within which the adaptive potential of activating *citT* expression changed due to one or more mutations. Specifically, we clustered phylogenetically-adjacent strains into groups within which all pairwise comparisons of the fitness effect of the P_*rnk*_*-citT* module were not significantly different (Bonferroni-corrected two-tailed bootstrap tests, *P* > 0.05). Overall, this analysis suggests that there were at least three major step-like changes in the potential for evolving the rudimentary Cit^+^ trait along the pre-Cit^‒^ lineage that eventually evolved citrate utilization (**Fig. 4**).

**Figure 3.**
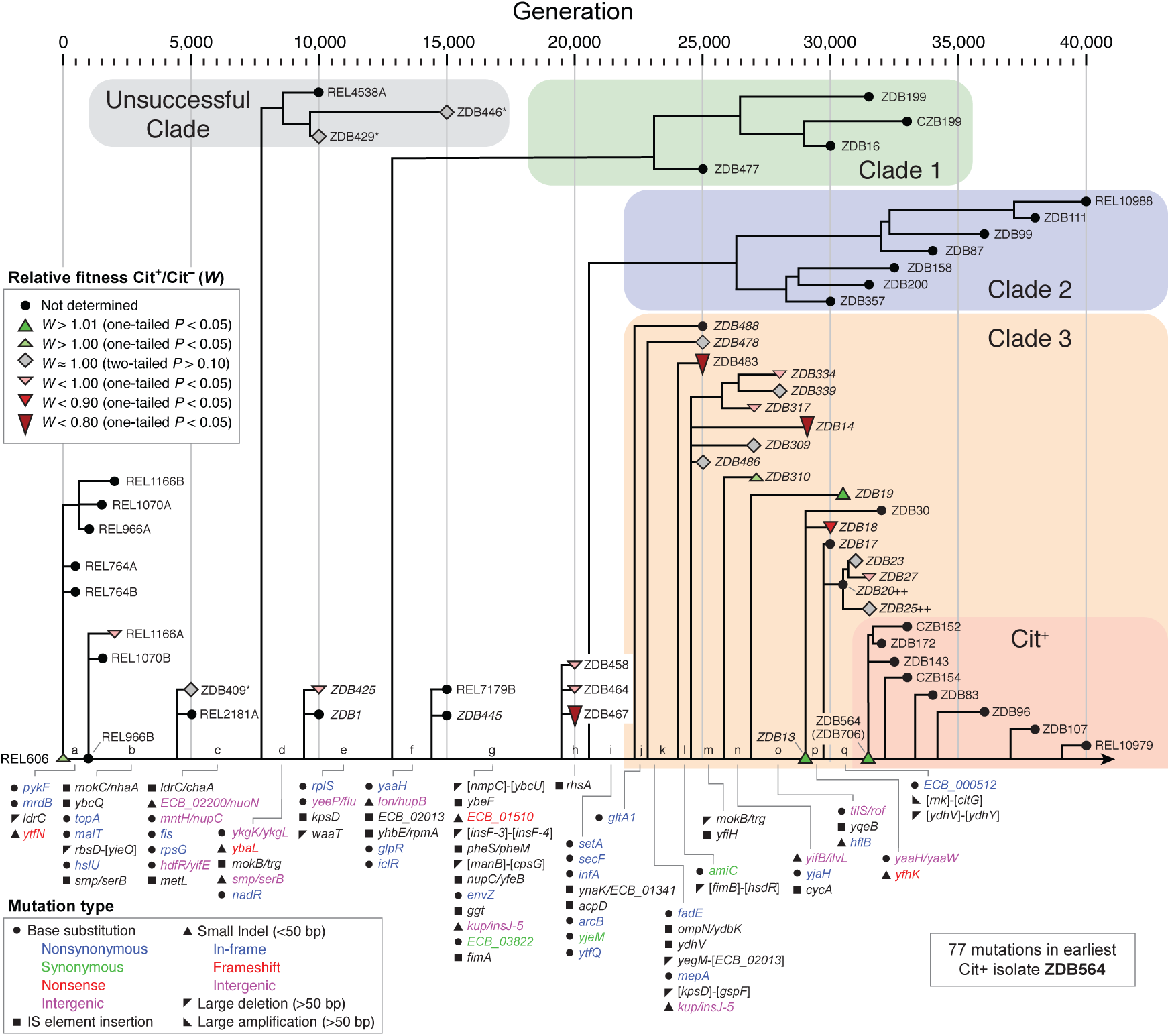
Potential for evolving Cit^+^ mapped onto phylogeny. Phylogeny of isolates from the LTEE population including 20 new clones sequenced for this study to provide better resolution of the timing of mutations on the lineage leading to Cit^+^ (names in italics). In order to identify changes in the degree of potentiation due to mutations, we mapped the results of the P_*rnk*_*-citT* knock-in assay onto this phylogenetic tree. Colored symbols reflect the Cit^+^ to Cit^‒^ relative fitness measured for those strains. The ancestor and 61 evolved isolates were used to construct this phylogenetic tree (**Table S1**). Two clones isolated at 50,000 generations are not shown. Two strains that evolved citrate utilization in replay experiments under the LTEE conditions in a previous study^7^ are marked with plus signs (++), and three strains that had evolved alleles added or removed during strain construction as described in **Table S2** are starred (*).

**Figure 4.**
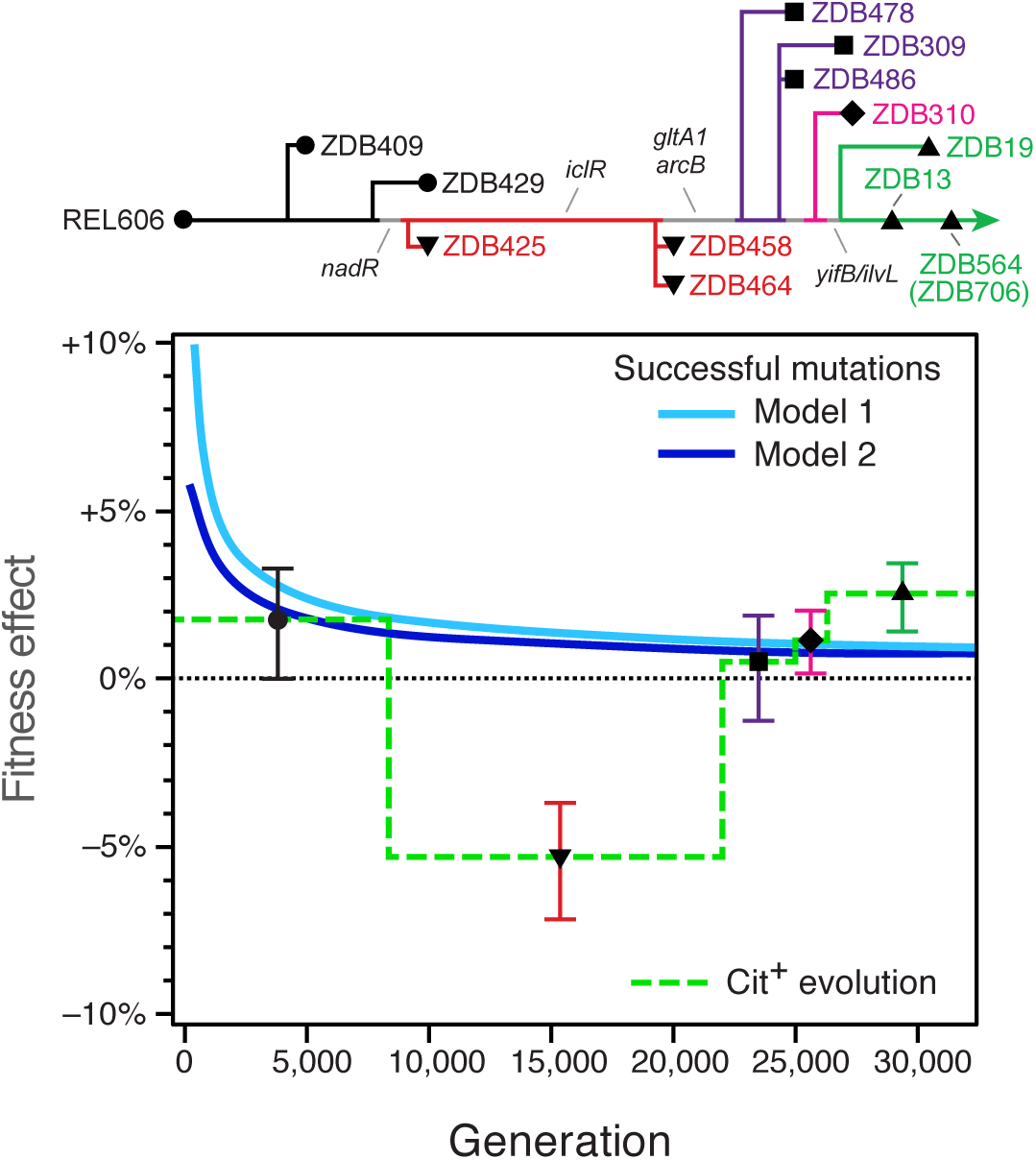
Changes in the potential for innovation along the lineage leading to Cit^+^ due to genetic and population factors. We clustered phylogenetically adjacent strains in which activating *citT* expression had a similar effect on fitness to reconstruct when major changes in potentiation occurred due to new mutations accumulating in the evolved strain (the genetic background). Each cluster is represented by a different color and symbol in the simplified phylogenetic tree (upper panel) and the graph showing the group-wise fitness estimate over a time period (horizontal lines) in which these strains were representative of the main pre-Cit^+^ lineage (lower panel). Error bars are 95% confidence intervals. Two models of the rate of adaptation of the LTEE populations at different generations are superimposed on the lower panel. Model 1 estimates the fitness effects of winning cohorts of beneficial mutations sequentially sweeping through the population at each generation according to modelling from Wiser et al.^14^. Model 2 estimates the fitness effects of each consecutive beneficial mutation accrued by the winning lineage over time using additional information from Tenaillon et al.^13^ (see **Methods**). These curves represent competing beneficial mutations that can suppress the evolution of Cit^+^ (the population context). If the fitness effect of activating *citT* is above these curves, as it is by 29,000 generations, then it is predicted to be among the most beneficial new mutations that could appear at that time in the LTEE. This means that rudimentary citrate utilization (Cit^+^) can persist in the population long enough to be refined by further mutations to full citrate utilization (Cit^++^); the metabolic innovation can be achieved.

Proceeding backward in the tree from the earliest known Cit^+^ isolate (ZDB564), two earlier clones (ZDB19 and ZDB13) from as early as 29,000 generations are as fully potentiated as the key Cit^‒^ revertant (ZDB706). The overall fitness effect of evolving Cit^+^ in this group was ^+^2.4% [^+^1.4%, ^+^3.4%] (95% confidence interval). The next-earliest group comprises three clones isolated at time points from 25,000 to 27,000 generations (ZDB478, ZDB486, and ZDB309). Activation of *citT* had little to no impact on this set of strains, with an estimated group-wise effect on fitness of ^+^0.4% [–1.3%, 2.0%]. One intermediate strain, ZDB310 from 27,000 generations, was not significantly different from either of these two groups immediately before and afterward, although the two groups were significantly different from one another.

It was deleterious to evolve Cit^+^ in an earlier, intermediate set of isolates composed of ZDB425, ZDB458, and ZDB464 with an estimated fitness effect of –5.4% [–7.2%, –3.6%]. These clones appear to be genetically typical of the pre-Cit^+^ lineage. ZDB425 at 10,000 generations and ZDB458 at 20,000 generations have only one and two ‘private’ mutations not shared with the main pre-Cit^+^ lineage, respectively, though we cannot rule out that other changes in the impact of *citT* activation may have occurred on the main line of descent within this interval. Before these anti-potentiated clones, there is an initial cluster that groups ZDB409 and ZDB429 with the REL606 ancestor. In these three isolates, evolution of Cit^+^ would have been slightly beneficial with a fitness impact of+1.7% [+0.0%, +3.3%].

Other strains are not classified into these major groups. It is less likely that they are representative of how potentiation evolved in the lineage leading to Cit^+^. For example, the four most highly anti-potentiated clones (ZDB467, ZDB483, ZDB14, ZDB18) appear to have evolved this property independently and due to ‘private’ mutations not shared with the main pre-Cit^+^ lineage (**Fig. 3**), at least this is the most parsimonious explanation. Similarly, the fitness effects measured in the P_*rnk*_*-citT* knock-in assay for two three-member subclades (ZDB334, ZDB339, ZDB317; and ZDB23, ZDB27, ZDB25) indicate that each likely shared one or more mutations that altered Cit^+^ potentiation only within that subclade, though the effects are much smaller in these cases. Finally, we excluded ZDB446 from this analysis because it was so deeply branched: removed by >5,000 generations from the pre-Cit^+^ lineage. It would have been clustered with the earliest group containing REL606 according to our criteria.

### Cit^+^ evolution in the context of competition with other beneficial mutations

During the time period when the pre-Cit^+^ lineage was anti-potentiated, from approximately 10,0 to 20,000 generations, invasion of a new Cit^+^ subpopulation would have been nearly impossible. Lineages that lost fitness by evolving the rudimentary version of this new trait would be rapidly purged by selection before refining mutations (e.g., activating *dctA)* could accumulate to give the decisive benefit of full citrate utilization (the Cit^++^ phenotype). What about the earlier and later time periods when the evolution of Cit^+^ was neutral or slightly beneficial? During these epochs, a newly evolved Cit^+^ lineage would still have had to compete with not only its own ancestor, but also against other lineages that were evolving at the same time, many of which would have other beneficial mutations. That is, an incipient Cit^+^ lineage had to survive in competition with alternative adaptive pathways, such as those improving fitness on glucose.

In order to understand when the fitness effects we measured for evolving Cit^+^ by *citT* activation would have made this metabolic innovation a viable evolutionary pathway in the context of competition within the LTEE population, we compared the group-wise fitness effects determined from the P_*rnk*_*-citT* knock-in assays to two models of the fitness effects of beneficial mutations that were successful at different generations in this LTEE population (**Fig. 4**). The Wiser *et al*. approach fits the fitness trajectory of this LTEE population to a model that incorporates a uniform type of diminishing returns epistasis between beneficial mutations and assumes consecutive sweeps^14^. The Tenaillon *et al*. model fits the number of beneficial mutations accumulating over time from genome sequencing data^13^. We combined this information with the Wiser *et al*. fitness trajectory to infer the representative fitness change for each subsequent beneficial mutation. The larger fitness effects in the Wiser *et al*. model reflect that it estimates the advantages of sweeping cohorts that may include more than one beneficial mutation. Overall, both models give very similar results that reflect the well-known deceleration in fitness gains during the Lenski long-term experiment^3,4,15,16^.

The models demonstrate that even if Cit^+^ evolution was marginally beneficial in the REL606 ancestor and other early isolates, it was initially much less beneficial than was needed to be successful at this point. Even by 5,000 generations, *citT* activation appears to have been average, at best, in terms of its fitness effect among all possible beneficial mutations. It would have been unlikely for the Cit^+^ trait to appear and persist at this point because there were so many alternative mutations, such as those that required only single-base substitutions or IS insertions that knocked out gene function, which would have occurred at a higher rate than the specific duplications or IS element insertions needed to activate *citT* expression^7^. After anti-potentiation appeared and receded in this lineage, competition would have continued to suppress Cit^+^ evolution when the *citT* mutation was again neutral. In striking contrast, evolving Cit^+^ was clearly superior to a typical successful beneficial mutation in the final group of strains that first evolved by 29,000 generations. It was a viable adaptive pathway at this point. Thus, by comparing P_*rnk*_*-citT* knock-in assays to models of the rates of population evolution, we can explain how a variant with a rudimentary Cit^+^ trait was able to appear and avoid extinction long enough to achieve the decisive *dctA* mutation that led to the dominant Cit^++^ trait.

## Discussion

Our work reframes and further elucidates why the emergence of citrate utilization is so rare in the Lenski long-term evolution experiment (LTEE). Rudimentary citrate utilization (the Cit^+^ phenotype) can apparently evolve at any time when a mutation switches on expression of the CitT transporter under the aerobic conditions of the experiment. However, the success of a new Cit^+^ variant is far from guaranteed. It is contingent on whether its descendants can survive long enough to incorporate a second mutation, such as one activating expression of the DctA transporter, that enables full citrate utilization (the Cit^++^ phenotype). The chance that Cit^++^ will be realized by this evolutionary pathway is dependent on two major factors. First, the initial mutational step conferring the weak Cit^+^ phenotype must be beneficial to fitness. Whether it is advantageous or not depends on the context of other mutations present in an evolved genome in which *citT* activation occurs. Second, the benefit of the mutation conferring weak Cit^+^ must be great enough that it can survive in competition with other adaptive mutations. Whether it is sufficiently beneficial depends on the population context in which it arises. We found that both genetic and population factors limited Cit^++^ evolution at different times in the LTEE (**Fig. 4**).

Unexpectedly, evolution of Cit^+^ by activating *citT* expression appears to have already been slightly beneficial to fitness in the ancestral strain used to found this *E. coli* population on the first day of the LTEE and to have remained so in other early evolved isolates. Even though Cit^+^ strains that evolved in the LTEE population at this point would have been capable of displacing their own Cit^‒^ ancestors, this first step on the pathway to the full Cit^++^ innovation was suppressed due to competition with mutations on adaptive pathways that improve fitness in the original glucose niche. New cells with highly beneficial mutations related to this primary component of the LTEE environment were essentially guaranteed to arise in the population and outcompete any cells with mutations activating *citT* expression. By 10,000 generations, the lineage in which Cit^+^ eventually evolved became ‘anti-potentiated’ after it accumulated additional mutations.

Now, the pathway to innovation was blocked because it was deleterious to evolve rudimentary Cit^+^ in this genetic background. There was a fitness valley separating the evolved Cit^‒^ strains from the full Cit^++^ phenotype. Finally, further mutations appeared in the focal LTEE lineage by 29,000 generations that altered the fitness impact of activating *citT* expression such that it was again beneficial to evolve the Cit^+^ phenotype, and perhaps even more so than it had been in the ancestor. At this point, the rate of adaptation of the population had slowed enough that evolving rudimentary Cit^+^ was now among the most beneficial mutational steps remaining. The two-step mutational pathway to Cit^++^ was no longer suppressed by genetic or population factors, and the Cit^++^ innovation evolved.

Cit^++^ mutants of *E. coli* capable of growth on citrate as a sole carbon source under aerobic conditions have been isolated in other studies^7,8,17,18^. In all of these cases, multiple mutations have been required to achieve the Cit^++^ phenotype. When they have been identified, the mutations that yield Cit^++^ activate expression of the CitT and DctA transporters, as is observed in the LTEE. These studies have isolated Cit^++^ mutants in much shorter periods of time (<1-8 weeks) than it took to evolve in the LTEE (~15 years) because they involve starving *E. coli* cells for days to weeks under conditions in which citrate was present as a potential carbon source. In the context of our results and as previously noted by others^19^, this difference in environmental conditions relative to the glucose-limited transfer regime of the LTEE, in which cells are in stationary phase for only ~16-18 hours each day, dramatically increases the fitness benefit of evolving the rudimentary Cit^+^ phenotype. Therefore, these stark conditions are expected to completely mask and overwhelm the dependency on potentiating genetic and population factors found in the LTEE. Activating *citT* expression would be universally beneficial in any genetic background in these types of experiments. Any increase in lag phase or other trade-off with respect to growth rate that might accompany this intermediate step in the pathway to Cit^++^ is irrelevant when cells without the mutation simply cannot replicate at all. The citrate-only starvation conditions also eliminate any interference from alternative mutations with benefits related to glucose utilization that suppress Cit^++^ evolution in the LTEE.

Why is evolution of Cit^+^ beneficial in some evolved genetic backgrounds and deleterious in others under the conditions of the LTEE? Activation of CitT expression under these aerobic conditions via the *rnk-citG* duplication leads to coupled import of citrate (a C_6_-tricarboxyate) and export of C_4_-dicarboxylates (e.g., succinate)^20^. In wild-type *E. coli* strains, CitT is normally expressed only under anaerobic conditions, and the imported citrate can only be assimilated when a fermentable co-substrate, such as glucose, is also present^21^. Under these conditions, citrate is cleaved to acetate and oxaloacetate by citrate lyase. The structural proteins and accessory factors necessary for producing this enzyme complex are encoded in the same operon as *citT*. When glucose is co-utilized with citrate, the resulting oxaloacetate is reduced to succinate by reverse tricarboxylic acid (TCA) cycle reactions. This process consumes reduced cofactors produced by breakdown of the sugar to balance redox metabolism without the need for O_2_. The succinate or other C_4_-dicarboxylates produced can be exchanged for more citrate import via CitT to continue this mixed fermentation mode of growth, or these TCA cycle intermediates can be siphoned off into biosynthetic pathways as necessary for cellular replication.

Under the aerobic conditions of the LTEE, citrate lyase is not expressed and succinate to balance citrate import by CitT must be produced in a different manner, from citrate or glucose using reactions of central metabolism. The availability of O_2_ makes it possible to maintain redox balance while synthesizing succinate via the TCA cycle, the glyoxylate bypass, or anaplerotic reactions (e.g., phosphoenolpyruvate carboxylase). *E. coli* growing under aerobic conditions ferments glucose to acetate, and mutations in genes related to the ability to re-uptake and utilize acetate are widespread in the LTEE^13,22,23^. These mutations affect acetate transporters and also pathways for assimilating acetate as acetyl-CoA through citrate synthase, the TCA cycle, and the glyoxylate bypass. Therefore, how these pathways are altered by adaptation to better utilize glucose and acetate is likely an important determinant of the genetic background that affects the ability to evolve citrate utilization. If introduction of the CitT transport reaction misbalances the redox state of the cell or the distribution of carbon compound intermediates between anabolism and catabolism, then it would be deleterious to fitness. Therefore, mutations altering central metabolism are candidates for explaining the changes in the fitness effect of *citT* activation along the LTEE lineage that ultimately evolved citrate utilization (**Fig. 4**).

Starting with the ancestor and examining when changes in the potential for evolving Cit^+^ were observed in the LTEE, a mutation in *nadR*, a repressor of NAD coenzyme biosynthesis^24^, occurs along the branch in the phylogenetic tree when anti-potentiation first evolved, before 10,000 generations. Mutations in *nadR* have appeared and swept to fixation in all twelve LTEE populations. These mutations include frameshift mutations and IS element insertions^13^, indicating that they are loss-of-function mutations, and deleting this gene from the genome of the LTEE ancestor has been shown to be beneficial^15^. Reducing or eliminating NadR activity is predicted to increase the NAD/NADH pool in the cell and could enable increased rates of glucose fermentation. Since NADH is a potent allosteric regulator of enzymes in central metabolism, including citrate synthase (*gltA*) for entry into the TCA cycle, this mutation may also reconfigure other cellular fluxes in ways that make CitT transport deleterious to fitness.

Between 10,000 and 25,000 generations mutations occurred in this LTEE population in three key genes that affected the activities of enzymes in central metabolism: *iclR, arcB*, and *gltA1*. These mutations have all been shown to improve growth on acetate^10^. Two of these mutations are in negative regulators; they are expected to derepress enzymes of the glyoxylate bypass (iclR)^25^ and TCA cycle (arcB)^26^, leading to increased metabolic flux through these pathways. Mutations in both of these genes are found in nearly all LTEE populations^13^. The citrate synthase mutation (*gltA1)* reduces allosteric inhibition of this enzyme by NADH^10^, which increases flux of acetyl-CoA into the TCA cycle. Evolved strains with mutations in *gltA* have only persisted in one other LTEE population that maintained the low ancestral mutation rate through 50,000 generations^13^. Both the *arcB* and *gltA1* mutations occurred on a branch in the phylogenetic tree for the citrate LTEE population when the effect of *citT* activation reverted to being neutral with respect to competitive fitness, so they are candidates for reversing anti-potentiation. The *iclR* mutation does not seem to have had an effect on genetic potentiation on its own, but it may have interacted with the *arcB* and/or *gltA1* mutations in a way that contributes to this anti-potentiation effect.

Only one mutation in a gene known to be involved in central metabolism occurred around 27,000 to 29,000 generations, at the point in the phylogenetic tree when adding the *citT* mutation seems to have again become beneficial to fitness. This mutation is upstream of the *ilv* operon for branched chain amino acid biosynthesis in the *yifB/ilvL* intergenic region. This pathway consumes pyruvate and acetyl-CoA, and its products can be used to synthesize the pantothenate moiety of coenzyme A (CoA)^27^. If this mutation affects gene expression of the *ilv* operon, then it could impact the balance of citric acid cycle intermediates flowing into or out of the TCA cycle to sustain cellular growth directly or indirectly via changing CoA/acetyl-CoA availability.

While the functions of the genes that we have highlighted in central metabolism suggest that they may be especially important for altering the potential for Cit^+^ evolution, other mutations also accumulated on the branches in the phylogenetic tree where the effects of *citT* activation on *E. coli* fitness changed (**Fig. 3**). In future work, the P_*rnk*_*-citT* knock-in assay can be used to further dissect this adaptive pathway by testing strains in which various evolved alleles have been removed or added. As an example of this type of approach, we have previously shown that removing the *gltA1* mutation from the earliest Cit^+^ isolate (ZDB564) makes the citT-activating duplication highly deleterious because it introduces a growth lag like that observed in the strongly anti-potentiated LTEE isolates in this study^10^. Similar studies could be conducted on strains that represent as closely as possible the genotypes present at critical junctures in the phylogenetic tree to determine which mutations altered the chances of achieving this innovation.

Another remaining question is whether the Cit^+^ innovation will ever evolve in the other eleven LTEE populations. It has not as of more than 60,000 generations^23^, nearly twice the amount of time that was required for it to evolve in the population analyzed here^7^. The ‘innovation interference’ of other highly beneficial mutations within a population suppressing Cit^+^ evolution has undoubtedly faded in all eleven of these populations as the pace of fitness increase has slowed similarly in all of them^14,28^. However, the ubiquity of *nadR* mutations in the LTEE may indicate that other populations similarly descended into a genetically anti-potentiated state. Our results suggest that Cit^++^ may still appear in the future if mutations suitably adjust fluxes in central metabolism to make evolving rudimentary Cit^+^ by activating *citT* expression a beneficial step on the pathway to innovation, as long as no critical components have been irrecoverably lost from the genome. Through 50,000 generations, no population has deleted either *citT* or *dctA*, and these genes have not accumulated any mutations in most populations^13^, so the latent genetic potential to evolve Cit^+^ seems to have remained intact so far.

The LTEE is an open-ended evolution experiment^29^; it did not begin with the aim of isolating *E. coli* that utilize citrate. There was never strong selection for this novel capability. Because evolving citrate utilization allowed the new Cit^++^ clade to colonize an untapped nutrient niche and rapidly diversify, this new metabolic capacity is an example of a key evolutionary innovation^30^. The evolution of Cit^++^ initiated a new round of rapid evolutionary optimization that included mutations that reduced the activity of citrate synthase (*gltA2)* and eliminated flux through the glyoxylate shunt (*aceA)*, both of which reversed the effects of pre-Cit^+^ adaptive mutations^10^. The many new possibilities for improving fitness in this alternative niche also likely contributed to the evolution of hypermutation within the Cit^++^ clade by 36,000 generations^8^. Lastly, new ecological interactions arose in this population such that Cit^‒^ and Cit^++^ types coexisted via negative-frequency dependent interactions for at least 10,000 generations after Cit^++^ evolved^7,8^. Continuing evolution of interactions between these and other *E. coli* lineages led to the emergence of an ecology that is unique to this flask in the LTEE^11^.

We found that a metabolic innovation in a laboratory population of *E. coli* was contingent on both a history of genetic adaptation and ongoing population dynamics. Evolution of metabolic capabilities has been found to be crucial to the emergence and continued success of bacterial pathogens in several instances^31,32^. For example, *Salmonella* acquired the ability to use tetrathionate as an electron acceptor, giving it a growth advantage relative to other bacteria in the environment that it creates in the gut during infection by inducing inflammation^33^. On a shorter timescale, mutations in the opportunistic pathogen *Pseudomonas aeruginosa* that accumulate during chronic infections in the cystic fibrosis lung lead to an increased ability to acquire iron from hemoglobin^34^. Even in the simple environment of the LTEE, both genetic and population factors suppress the evolution of an innovation that allows a new niche to be exploited by a new bacterial species. It may be useful in the treatment of disease to understand when these and other factors, including competition for specific nutrients by commensal species in a microbiome, can be used to suppress evolutionary outcomes that are harmful to human health^35^.

## Materials and Methods

### Media conditions and strains

*E. coli* were cultured in Davis-Mingioli (DM) medium and Lysogeny Broth (LB)^10^. As necessary, media were supplemented with 50 μg/mL kanamycin and 80 μg/mL 5-bromo-4-chloro-3-indolyl β-D-galactopyranoside (X-gal). Evolved clones characterized in this study from archived LTEE populations and strain ZDB706 (the spontaneous Cit^‒^ revertant of ZDB564) were isolated in previous studies^7,8,10^. New strains constructed in this study are listed in **Table S3**.

### P_*rnk*_*-citT* knock-in assay

The activated P_*rnk*_*-citT* module was constructed by amplifying the evolved *rnk-citG* duplication junction from the pCit plasmid along with a linked kanamycin resistance gene (Kan^r^)^9^. The P_*rnk*_*-citT* construct in pCit is originally from evolved strain CZB154^9^. As a control, another module was created which only contains the Kan^r^ marker. These modules were integrated into the genomes of several Cit^‒^ strains (REL607, REL1166A, ZDB429, ZDB467, and ZDB483) via lambda Red recombination^36^ such that they replaced the *lac* locus (*lacA* to *lacZ)*, spanning positions 333,862-337,485 in the REL606 genome (GenBank:NC_012967.1)^37^. We transferred the cassettes to other strains using P1 bacteriophage transduction^38^. Successful transductants were scored based on blue/white screening in the presence of X-gal and kanamycin. All Cit^+^ strains were made by transduction of the P_*rnk*_*-citT* module into an Ara^‒^ LTEE clone. Isogenic Cit^‒^ strains were constructed by insertion of the control Kan^r^ module into an Ara^+^ version of the same clone generated as described in the next section. To determine whether any other mutations present in the evolved strains from the LTEE were altered during transduction, we screened for mutations identified by whole-genome sequencing in the recipient strain that were within 100 kb upstream or downstream of the P_rnk_-*citT* insertion site. Strains from three Cit^+^/Cit^‒^ pairs were found to have gained or lost evolved alleles in this process (**Table S2**).

### Selection for spontaneous Ara^+^ mutants

All Ara^‒^ strains inherited a point mutation in *araA* present in the REL606 LTEE ancestor that prevents arabinose utilization^15^. To isolate spontaneous Ara^+^ mutants, Ara^‒^ strains were revived overnight at 37°C in DM containing 1 mg/mL glucose (DM1000). For each strain, three separate flasks containing 10 ml of DM1000 were each inoculated with ~500 cells from the first DM1000 culture to reduce the chance that they might share any secondary mutations affecting fitness. After incubating overnight at 37°C, cells were harvested by centrifugation at 4,000 rpm for 15 min and the entire volume was plated on minimal arabinose (MA) plates. Plates were incubated for 36-48 h and colonies were streaked and grown on new MA plates before picking single-colonies as candidate Ara^+^ revertants. The presence of secondary mutations affecting fitness was assessed by competing the original Ara^‒^ and selected Ara^+^ strains, as described below. In most cases, we identified an Ara^+^ revertant with a fitness that was not significantly different from its Ara^‒^ progenitor (**Fig. S3**).

### Competition assays

Relative fitness was measured using co-culture competition assays^4,39^. Two strains to be competed are differentiated based on their ability to ferment arabinose. Ara^‒^ strains form red colonies on tetrazolium arabinose (TA) media, and Ara^+^ strains form pink colonies. Strains were revived overnight in LB then were diluted 10,000-fold into separate cultures for each replicate competition assay in DM containing 25 μg/mL glucose (DM25).

These cultures were preconditioned and competed under the same conditions as used in the LTEE^4,40^, in 10 mL of DM25 in 50 mL Erlenmeyer flasks shaken at 120 rpm over a diameter of 1 inch with incubation at 37°C. After 24 h of growth separately to precondition strains to these conditions, two replicate cultures for each Ara^‒^ and Ara^+^ pair were mixed at equal volumes in fresh DM25 media such that there was an overall 1:100 dilution. Dilutions of these initial mixtures were plated on TA plates to determine the initial representation of each strain in each replicate flask. Then, the competition was carried out over three days of transferring 1:100 dilutions into fresh medium each day. A dilution of each culture after growth on day three was again plated to determine the final representation of each strain. Relative fitness was calculated as the ratio of the realized growth rates of each strain between the final and initial platings^4,39^.

For comparisons of the effect of the authentic *rnk-citG* duplication versus the addition of the P_*rnk*_*-citT* module to REL606 and ZDB706 (**Fig. 1**) we first established neutrality of an Ara^+^ revertant and then judged whether there was significant difference between the fitnesses of the Cit^‒^ and Cit^+^ strains pairs. For comparing the fitness impact of evolving Cit^+^ in other strains (**Fig. 2**), we measured the relative fitness of the Ara^‒^ Cit^+^ variant of the strain with the P_*rnk*_*-citT* module added versus the Ara^+^ Cit^‒^ revertant of its Cit^‒^ progenitor (Cit competition) and multiplied this by the relative fitness of the Ara^+^ Cit^‒^ revertant versus the Ara^‒^ Cit^‒^ clone with the null module added (Ara competition) (**Fig. S2**). To account for how error in each of these two competitions impacts confidence in the overall fitness change inferred for evolving Cit^+^, we performed 10,000 bootstrap resamplings of the Ara and Cit competition replicates to estimate 95% fitness intervals and significance on the combined measurements. The same bootstrapping procedure was used for comparing the fitnesses of different strains in the population phylogeny in the procedure that combined them into equivalence groups along the lineage to Cit^+^ (Fig. 4).

qRT-PCR measurement of *citT* expression. Cells were cultured according to the method described in Blount *et* al.^8^. Briefly, cells were initially grown to saturation in a 5 ml LB culture and transferred into 10 ml of DM25 media (1:10,000 dilution) followed by two 24-hour preconditioning cycles in DM25 with 1:100 dilutions. For each preconditioning cycle, cells were diluted by 1:100 into fresh DM25 media. At this point, we performed a final dilution of 1:100 into DM25. Cells were grown until they reached ~50% of the final OD_420_ and the entire culture (10 ml) was harvested for extracting RNA. RNA was extracted from frozen cell pellets using the RNASnap protocol^41^. The resulting supernatant was column purified, incorporating on-column DNase treatment (RNA Clean & Concentrator-25, Zymo Research). TapeStation analysis (Agilent) was used to verify RNA integrity (all RIN scores ≥ 8.0). Samples were then reverse transcribed in parallel using random primers, with 200ng of RNA as template (High Capacity Reverse Transcription Kit, Applied Biosystems).

qPCR was run in 384 well plates on an Applied Biosystems ViiA 7, using SYBR Green (Thermo Fisher) as fluorophore in a 5 μl reaction. QuantStudio was used to determine quantification cycle (Cq) values. All samples were run in technical triplicates. We selected two reference genes (*refs)*, 16S RNA and *idnT*, from an initial pool of candidates based on primer efficiency, primer specificity (as judged by melt curve) and stability of expression in a subset of our strains of interest. Primer efficiency was calculated from the slope of a plot of log(dilution) versus Cq, using a 5-fold or 10-fold dilution series of a pool of cDNA from every sample. Final primer sequences and efficiencies were as follows: *citT* (forward = GTTATAGCGGGTAATGTCTTTC, reverse = CACTGATTGGCCTTGTATTG, efficiency = 99.25%); *idnT* (forward = CCCGACACCGCTATCTACTAATAC, reverse = CGCACCATCGAGCAAATCAT, efficiency = 100.5%); 16S (forward = CCCGAAGGTTAAGCTACCTACT, reverse = CATGAAGTCGGAATCGCTAGTAATC, efficiency = 97.6%). In our final analysis comparing *citT* expression across strains, we used three biological replicates per strain, and 2 μl of a 1:100 dilution of cDNA as template. Relative expression (R) of *citT* in the strain of interest relative to ancestral REL606 was calculated as follows. First, DCq was calculated for individual biological replicates according to ΔCq = Cq^*citT*^ – *x̄*(Cq^*refs*^), where *x̄*(*X*) represents the mean of the values for quantity *X*. Then, ΔΔCq and *R* were calculated from the mean ΔCq of three biological replicates for each strain tested as ΔΔCq = *x̄*(ΔCq^strain^) – *x̄*(ΔCq^REL606^) and *R* = 2^-ΔΔCq^.

### Genome sequencing and phylogenetic tree construction

All 61 evolved strains from the LTEE population with genomes analyzed in this study are listed in **Table S1**. For the 20 newly sequenced strains, genomic DNA was purified using the GenElute Bacterial Genomic DNA kit (Sigma) and then sequenced using standard procedures on an Illumina HiSeq 2500 instrument to generate 101-base paired-end reads by the University of Texas at Austin Genome Sequencing and Analysis Facility. Data files for these 20 genomes have been deposited in the NCBI Sequence Read Archive (SRP120037). Raw sequencing reads for all 61 genomes are available via links from the main LTEE NCBI BioProject page (PRJNA414462).

We initially predicted mutations in each re-sequenced genome by comparing Illumina reads to the REL606 reference genome^37^ using *breseq* (v0.31.1)^42,43^. Then, we further curated the lists of predicted mutations as previously described^13^. Briefly, a maximum-parsimony phylogenetic tree for all 61 strains from the LTEE population was constructed using the DNAPARS program from the PHYLIP package (v3.69)^44^. Where necessary, we manually corrected mutation predictions, including adding mutations that were hidden by later deletions or splitting sequence differences into multiple mutational events to construct the most parsimonious phylogeny possible. In the current study, we did not discard mutations in repetitive regions before analysis, except we did ignore changes in the hypervariable 7×CCAG repeat at reference coordinates 2103891-2103918. The final lists of mutations predicted in all clones are available online in version 2.0 of the GitHub LTEE-Ecoli repository (https://github.com/barricklab/LTEE-Ecoli).

### Beneficial mutation fitness effect models

To construct the curve for the Wiser et al. model^14^ in Figure 4 we calculated the expected time in generations (*t*) and fitness increase (*s*) for each subsequent sweep of a cohort of beneficial mutations using equations S3, S4, and S7 from the supplement of that study using parameter values (*α*_*0*_ = 58.4, *μ* = 10^-7^, and *N* = 3.3×10^7^) that they found to be compatible with the fitness trajectories of the non-mutator LTEE populations. For the Tenaillon et al. model^13^, we first calculated a curve describing the number of beneficial mutations expected in an evolved isolate (*n*) according to the term, 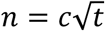 with the best-fit coefficient value (*c* = 0.135) found in that study for all non-mutator LTEE populations considered together. Next, we combined this model with the fitness (*W)* model from Wiser et al., *W*(*t*) = (*at* + 1)^*b*^, with best-fit parameters (*a* = 0.0842 and *b* = 0.00611) found specifically for the citrate population (Ara–3)^14^. Finally, the Tenaillon et al.^13^ curve in **Figure 4** was graphed by calculating each generation (*t*_*n*_) at which the number of beneficial mutations (*n*) was an integral value, *t*_*n*_ = (*n*/*c*)^2^. The graphed selection coefficient was estimated as the fitness at that time *W*(*t*_*n*_) minus the fitness at the time of the previous beneficial mutation *W*(*t*_*n-1*_).

## Acknowledgments

We thank Zachary Blount, Noah Ribeck, and Michael Wiser for helpful discussions and Craig Barnhart for assistance with genome sequencing. We acknowledge the Texas Advanced Computing Center (TACC) at The University of Texas at Austin for providing high-performance computing resources. This work was funded by the Army Research Office (W911NF-12-1-0390), National Institutes of Health (R00-GM087550), and the National Science Foundation BEACON Center for the Study of Evolution in Action (DBI-0939454). D.L. acknowledges a Robert D. Watkins Graduate Research Fellowship from the American Society for Microbiology.

## References

1. Davis, B. D. & Mingioli, E. S. Mutants of Escherichia coli requiring methionine or vitamin B12. J Bacteriol 60, 17‒28 (1950).

2. Blount, Z. D. A case study in evolutionary contingency. Stud Hist Philos Biol Biomed Sci 58, 82‒92 (2016).

3. Lenski, R. E. & Travisano, M. Dynamics of adaptation and diversification: a 10,000-generation experiment with bacterial populations. Proc Natl Acad Sci USA 91, 6806‒6814 (1994).

4. Lenski, R. E., Rose, M. R., Simpson, S. C. & Tadler, S.C. Long-term experimental evolution in Escherichia coli. I. Adaptation and divergence during 2,000 generations. The American Naturalist 138, 1315‒1341 (1991).

5. Rozen, D. E. & Lenski, R. E. Long-term experimental evolution in Escherichia coli. VIII. Dynamics of a balanced polymorphism. The American Naturalist 155, 24‒35 (2000).

6. Le Gac, M., Plucain, J., Hindre, T., Lenski R.E. & Schneider, D. Ecological and evolutionary dynamics of coexisting lineages during a long-term experiment with Escherichia coli. Proc Natl Acad Sci USA. 109, 9487‒9492 (2012).

7. Blount, Z. D., Borland, C. Z., Lenski, R. E. Historical contingency and the evolution of a key innovation in an experimental population of Escherichia coli. Proc Natl Acad Sci USA 105, 7899‒7906 (2008).

8. Blount, Z. D., Barrick, J. E., Davidson, C. J. & Lenski, R. E. Genomic analysis of a key innovation in an experimental Escherichia coli population. Nature 489, 513‒518 (2012).

9. Quandt, E. M., Deatherage, D. E., Ellington, A. D., Georgiou, G. & Barrick, J. E. Recursive genomewide recombination and sequencing reveals a key refinement step in the evolution of a metabolic innovation in Escherichia coli. Proc Natl Acad Sci USA 111, 2217‒2222 (2014).

10. Quandt, E. M., Gollihar, J., Blount, Z.D., Ellington, A.D., Georgiou, G., Barrick, J.E. Fine-tuning citrate synthase flux potentiates and refines metabolic innovation in the Lenski evolution experiment. eLife Sciences 4, e09696 (2015).

11. Turner, C. B., Blount, Z. D., Mitchell, D. H., Lenski, R.E. Evolution and coexistence in response to a key innovation in a long-term evolution experiment with Escherichia coli. bioRxiv 020958, doi: https://doi.org/10.1101/020958 (2015).

12. Barrick, J. E. & Lenski, R. E. Genome dynamics during experimental evolution. Nat Revs Genet 14, 827‒839 (2013).

13. Tenaillon, O., Barrick, J.E., Ribeck, N., Deatherage, D.E, Blanchard, J.L., Dasgupta, A., Wu, G.C., Wielgoss, S., Cruveiller, S., Medigue, C., Schneider, D. & Lenski R.E. Tempo and mode of genome evolution in a 50,000-generation experiment. Nature 536, 165‒170 (2016).

14. Wiser, M. J., Ribeck, N., Lenski, R. E. Long-term dynamics of adaptation in asexual populations. Science 342, 1364‒1367 (2013).

15. Barrick, J. E. Yu, D.S., Yoon, S.H., Jeong, H., Oh, T.K., Schneider, D., Lenski, R.E. & Kim, J.F. Genome evolution and adaptation in a long-term experiment with Escherichia coli. Nature 461, 1243‒1247 (2009).

16. Khan, A. I., Dinh, D. M., Schneider, D., Lenski, R. E., Cooper, T. F. Negative epistasis between beneficial mutations in an evolving bacterial population. Science 332, 1193‒1196 (2011).

17. Hall, B. G. Chromosomal mutation for citrate utilization by Escherichia coli K-12. J Bacteriol 151, 269‒273 (1982).

18. van Hofwegen, D. J., Hovde, C. J., Minnich, S. A. Rapid Evolution of Citrate Utilization by Escherichia coli by Direct Selection Requires citT and dctA. J Bacteriol 198, 1022‒1034 (2016).

19. Roth, J. R. & Maisnier-Patin S. Reinterpreting long-term evolution experiments: Is delayed adaptation an example of historical contingency or a consequence of intermittent selection? J Bacteriol 198, 1009‒1012 (2016).

20. Pos, K. M., Dimroth, P. & Bott, M. The Escherichia coli citrate carrier CitT: a member of a novel eubacterial transporter family related to the 2-oxoglutarate/malate translocator from spinach chloroplasts. J Bacteriol 180, 4160‒4165 (1998).

21. Lütgens, M. & Gottschalk, G. Why a co-substrate is required for anaerobic growth of Escherichia coli on citrate. J Gen Microbiol 119, 63‒70 (1980).

22. Barrick, J. E. & Lenski, R. E. Genome-wide mutational diversity in an evolving population of Escherichia coli. Cold Spring Harb Symp Quant Biol 74, 119‒129 (2009).

23. Good, B. H., McDonald, M. J., Barrick, J. E., Lenski, R. E. & Desai, M. M. The dynamics of molecular evolution over 60,000 generations. Nature 551, 45‒50 (2017).

24. Raffaelli, N. Lorenzi, T., Mariani, P.L., Emanuelli, M., Amici, A., Ruggieri, S. & Magni, G. The Escherichia coli NadR regulator is endowed with nicotinamide mononucleotide adenylyltransferase activity. J Bacteriol 181, 5509‒5511 (1999).

25. Cortay, J. C., Negre, D., Galinier, A., Duclos, B., Perriere, G. & Cozzone, A.J. Regulation of the acetate operon in Escherichia coli: purification and functional characterization of the IclR repressor. EMBO J. 10, 675‒679 (1991).

26. Perrenoud, A. & Sauer, U. Impact of global transcriptional regulation by ArcA, ArcB, Cra, Crp, Cya, Fnr, and Mlc on glucose catabolism in Escherichia coli. J Bacteriol 187, 3171‒3179 (2005).

27. Salmon, K. A., Yang, C.-R. & Hatfield, G. W. Biosynthesis and regulation of the branched-chain amino acids. EcoSal Plus doi:10.1128/ecosalplus.3.6.1.5 (2013).

28. Lenski, R. E., Wiser, M.J., Ribeck, N., Blount, Z.D., Nahum, J.R., Morris, J.J., Zaman, L., Turner, C.B., Wade, B.D., Maddamsetti, R., Burmeister, A.R, Baird, E.J., Bundy, J., Grant, N.A., Card, K.J., Rowles, M., Weatherspoon, K., Papoulis, S.E., Sullivan, R., Clark, C., Mulka J.S. & Hajela, N. Sustained fitness gains and variability in fitness trajectories in the long-term evolution experiment with Escherichia coli. Proc Biol Sci 282, 20152292 (2015).

29. Fox, J. W. & Lenski, R. E. From here to eternity—the theory and practice of a really long experiment. PLoS Biol 13, e1002185 (2015).

30. Heard, S. B. & Hauser, D.L. Key evolutionary innovations and their ecological mechanisms. Taylor & Francis 10, 151‒173 (1995).

31. Rohmer, L., Hocquet, D. & Miller, S. I. Are pathogenic bacteria just looking for food? Metabolism and microbial pathogenesis. Trends Microbiol 19, 341‒348 (2011).

32. Didelot, X., Walker, A. S., Peto, T. E., Crook, D. W., Wilson, D. J. Within-host evolution of bacterial pathogens. Nat Rev Microbiol 14, 150‒162 (2016).

33. Winter, S. E., Thiennimitr, P, Winter, M.G., Butler, B.P., Huseby, D.L., Crawford, R.W., Russell, J.M., Bevins, C.L, Adams, L.G., Tsolis, R.M., Roth, J.R. & Baumler, A.J. Gut inflammation provides a respiratory electron acceptor for Salmonella. Nature 467, 426‒429 (2010).

34. Marvig, R. L., Damkiær, S. Hossein Khademi, S.M., Markussen, T.M., Molin, S., Jelsbak, L. Within-host evolution of Pseudomonas aeruginosa reveals adaptation toward iron acquisition from hemoglobin. MBio 5, e00966‒14 (2014).

35. Bull, J. J. & Barrick, J. E. Arresting Evolution. Trends Genet 33, 910‒920 (2017).

36. Datsenko, K. A. & Wanner, B. L. One-step inactivation of chromosomal genes in Escherichia coli K-12 using PCR products. Proc Natl Acad Sci USA 97, 6640‒6645 (2000).

37. Jeong, H., Barbe, V., Lee, C.H., Vallenet, D., Yu, D.S., Choi, S.H., Couloux, A., Lee, S.W, Yoon, S.H., Cattolico, L., Hur, C.G., Park, H.S., Ségurens, B., Kim, S.C., Oh, T.K., Lenski, R.E., Studier, F.W., Daegelen, P. & Kim, J.F. Genome sequences of Escherichia coli B strains REL606 and BL21(DE3). J Mol Biol 394, 644‒652 (2009).

38. Thomason, L. C., Constantino, N., Court, D. L. E. coli genome manipulation by P1 transduction. Curr Protoc Mol Biol Chapter 1, Unit 1.17 (2007).

39. Wiser, M. J. & Lenski, R. E. A Comparison of Methods to Measure Fitness in Escherichia coli. PLoS ONE 10, e0126210‒11 (2015).

40. Lenski, R. E. Phenotypic and genomic evolution during a 20,000-generation experiment with the bacterium Escherichia coli, in Plant Breeding Reviews: Longterm Selection: Crops, Animals, and Bacteria, Volume 24, Part 2 (ed J. Janick), John Wiley & Sons, Inc., Oxford, UK. (2003).

41. Stead, M. B., Agrawal, A., Bowden, K. E., Nasir, R., Mohanty, B.K., Meagher, R.B. & Kushner, S.R. RNA snap ™: a rapid, quantitative and inexpensive, method for isolating total RNA from bacteria. Nucleic Acids Res 40, e156 (2012).

42. Deatherage, D. E. & Barrick, J. E. Identification of mutations in laboratory-evolved microbes from next-generation sequencing data using breseq. Methods Mol Biol 1151, 165‒188 (2014).

43. Barrick, J. E., Colburn, G., Deatherage, D.E., Traverse, C.C., Strand, M.D., Borges, J.J., Knoester, D.B., Reba, A. & Meyer, A.G. Identifying structural variation in haploid microbial genomes from short-read resequencing data using breseq. BMC Genomics 15, 1039 (2014).

44. Feisenstein, J. PHYLIP-Phylogeny Inference Package (Version 3.2). Cladistics 5, 164‒166 (1989).

